# Adaptive immunity shapes baseline physiology of *M. tuberculosis* in high-dose versus low-dose infection BALB/c mouse drug treatment models

**DOI:** 10.64898/2026.03.26.714401

**Authors:** Jo Hendrix, Reem Al Mubarak, Karen Rossmassler, Holly Nielsen, Elizabeth Wynn, Camile Moore, Isabelle L. Jones, Martin I. Voskuil, Brendan K. Podell, Gregory T. Robertson, Chuangqi Wang, Nicholas D. Walter

## Abstract

Preclinical tuberculosis (TB) drug evaluation relies heavily on mouse infection models. Drug efficacy varies depending on inoculum and the timing of treatment initiation. These differences reflect, in part, physiological adaptations of *Mycobacterium tuberculosis* (*Mtb*) to host immune pressure. We used novel molecular markers of pathogen health (RS ratio and SEARCH-TB) to contrast *Mtb* phenotypes in the BALB/c high dose aerosol (HDA) subacute TB and low dose aerosol (LDA) chronic TB infection models, focusing on the transition from innate to adaptive immunity. We found that the onset of adaptive immunity coincided with a rapid reprogramming of *Mtb* physiology, characterized by suppressed respiration, metabolism, and biosynthesis, together with induction of stress and nutrient acquisition pathways. Changes in key *Mtb* processes were concordant with changes in host expression of canonical features of adaptive immunity. Our results also explain key model-specific differences in drug efficacy: when used for drug evaluation, the HDA model begins treatment during the innate immune phase when *Mtb* are metabolically active, whereas the LDA model initiates therapy after host adaptive immunity has already activated and the bacterial population is immune-constrained. The HDA model is shown to be a mixed model, spanning immune phases with initial treatment acting on replicating bacteria and later treatment acting on immune-constrained populations. Together, the two models offer complementary perspectives on therapeutic activity across the spectrum of bacterial states and provide a framework for designing regimens that are effective against both active and immune-constrained *Mtb*.

## INTRODUCTION

Tuberculosis (TB) remains the leading cause of death from infection worldwide.^1^ Challenges include the need for prolonged treatment durations and drug resistance. There is an urgent need for optimized drug combinations that can cure all forms of TB more rapidly. Preclinical drug and combination drug regimen evaluation relies heavily on mouse models, but drug efficacy varies depending on mouse strain, infectious dose, and timing of treatment initiation.^2–5^ Because murine studies guide regimen selection for clinical trials,^2–5^ understanding model-specific differences is essential. Additionally, model differences may provide insights into drug activity that are relevant to human TB treatment. Here, we contrasted baseline *Mycobacterium tuberculosis* (*Mtb*) phenotypes in the absence of treatment in two widely used mouse infection models, the BALB/c high dose aerosol (HDA) and low dose aerosol (LDA) models,^5^ while evaluating the influence of adaptive immunity on *Mtb* physiology.

Several factors can contribute to variation in efficacy between murine models, including drug absorption, distribution, metabolism, and excretion^6–10^ and the physiologic state of *Mtb* at treatment initiation. Here, we focus on physiology. *Mtb* adapts to host immunity and diverse microenvironments varying in O_2_, CO_2_, pH, carbon and nitrogen sources, and micronutrients through physiological adaptations, including shifts in growth rate, metabolism, and stress responses. These adaptations are known to influence drug efficacy.^7,11–13^ Given the dependence of drug efficacy on *Mtb* physiological state, there is need for better understanding of the baseline pre-treatment *Mtb* phenotypes in the standard murine models used for regimen selection.

While both are conducted in the BALB/c mouse, the LDA and HDA models differ in inoculum and timing of treatment start. In the HDA model, aerosol deposition of a high bacterial load (∼10^4^ bacilli) in the lungs causes overwhelming lethal disease by day 20 unless mice are “rescued” by drug treatment. Treatment in the HDA model typically begins on day 11 or 14 post-aerosol while *Mtb* is actively replicating during the innate immune period. In the LDA model, deposition of a lower bacterial load (∼10^2^ bacilli) allows adaptive immune containment (activated around day 16),^14,15^ leading to chronic stable infection with slow bacterial replication.^16–18^ Treatment in the LDA model typically begins on day 28 or later, well after the onset of adaptive immunity. The HDA model is used in relapse studies to characterize the sterilizing activity of combination regimens.^19^ Because the LDA infection model results in immune-constrained, slowly replicating Mtb, it is often considered a more stringent drug efficacy model, and is therefore used for late stage evaluation of individual drugs.^5^

Certain aspects of *Mtb* physiology in the HDA and LDA models are known. In both models, exponential increase in colony forming units (CFU) during early infection reflects a rapidly growing *Mtb* phenotype.^14,20^ With the onset of adaptive immunity, transcriptional studies in LDA infections have shown that macrophage activation impairs aerobic respiration, causing metabolic shifts and slowed replication.^21–24^ In immunocompetent mice, CFU plateaus or declines slightly as adaptive immunity contains infection,^14,20^ though plasmid “replication clock” experiments indicate continued, albeit reduced, bacterial replication.^16–18^ However, critical gaps remain. First, most transcriptional studies were conducted in C57BL/6 mice^18,23^ which exhibit stronger Th1 polarization than BALB/c mice^2–5^ that are more commonly used for drug evaluation. Additionally, prior work focused on LDA infection alone and comparisons with HDA are lacking. Finally, most studies used qPCR^23–27^ (targeting few transcripts) or microarray^12,22,28–31^ (with limited sensitivity in mouse tissue), capturing only partial transcriptional information. Here, we used two novel molecular methods to probe pathogen health: (1) the RS ratio^®^ assay which quantifies *Mtb* activity based on ongoing rRNA synthesis^20^ and (2) SEARCH-TB, a targeted RNA-seq platform that amplifies *Mtb* mRNAs to quantify the *Mtb* transcriptome.^32^ As previously demonstrated,^32^ SEARCH-TB is substantially more sensitive than microarray and assays more targets than is feasible with qPCR, providing a highly granular measure of *Mtb* gene transcription. In the LDA model, a 56-day time course captured host and *Mtb* transcriptional changes from early infection through chronicity, with focus on shifts around adaptive immunity. In the HDA model, a 20-day time course highlighted *Mtb* responses under overwhelming infection. We also compared pre-treatment *Mtb* transcriptomes (day 11 in HDA vs. day 28 in LDA) to assess how dose and immune context shape bacterial state. These analyses clarified the impact of adaptive immunity within and across models.

In addition, we also performed RNA-seq on mice in the LDA model to investigate host-pathogen interactions during the onset of adaptive immunity. Previous studies generally evaluated either *Mtb*^12,13,23–31^ or the mouse.^15,33^ In cases where investigators evaluated transcriptional changes in both the host and the pathogen,^34,35^ the studies were limited to one time point and did not assess concordance between host and pathogen expression. To further investigate host-pathogen dynamics, we examined key host immune processes in relation to *Mtb* processes and identified concordance over time, thereby enhancing understanding of preclinical models that are central to contemporary drug development.

## METHODS

### Infection and sample collection

Animal procedures adhered to national and international guidelines under supervision of the Colorado State University Animal Care and Use Committee (Supplemental Information). For the chronic infection model, female mice (6-8 weeks old) were infected with a low dose aerosol of *Mtb* Erdman using the Glas-Col Inhalation Exposure System. Five mice each were sacrificed on infection days 1, 7, 11, 19, 28, and 56. An additional 15 mice were infected with a high dose aerosol of *Mtb* Erdman. Five mice were sacrificed on infection days 1, 11, and 19. Lungs were flash-frozen in liquid nitrogen for immediate RNA preservation.

### CFU Enumeration

Left lung lobes were homogenized in 4ml of 1×PBS using a Precellys tissue homogenizer then serially diluted in 1×PBS and plated on 7H11-OADC agar (i.e., Middlebrook 7H11 agar plates supplemented 0.2% [v:v] glycerol, 10% [v:v] oleic acid-albumin-dextrose-catalase (OADC) supplement, and 0.01 mg/mL cycloheximide, and 0.05 mg/mL carbenicillin). Plates were incubated at 37°C in a dry-air incubator for at least 21 days before counting. The bacterial doubling time was calculated as follows:

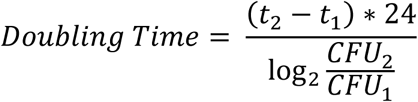

where CFU_1_ represents the mean CFU at time t_1_, selected randomly with replacement from available samples at t_1_. CFU_2_ was determined similarly for t_2_. A 95% confidence interval was estimated by recalculating the doubling time using 1000 rounds of bootstrapping with the boot^36^ function in R, followed by confidence interval determination using the boot.ci function from the boot package.

### RNA extraction and Pathogen and Host Sequencing

Right upper lung lobes were thawed in a 4M guanidine thiocyonate solution and homogenized (Precellys Evolution, Bertin) before lysis of *Mtb* via bead-beating with 0.1 zirconium oxide beads (CK01 tubes). RNA was purified (Maxwell RSC, Promega) simplyRNA tissue kit and prepared for *Mtb-*targeted RNA-seq (SEARCH-TB) using Illumina’s AmpliSeq for Illumina Custom and Community RNA Panels kit. Libraries were sequenced using Illumina NovaSeq6000. Additional information is detailed in Supplemental Methods.

For host sequencing, we prepared host libraries with the Universal Plus mRNA-Seq kit (Tecan) and generated paired-end sequencing reads of 150bp on NovaSeq 6000 (Illumina) sequencer at a target depth of 80 million paired-end reads per sample. Raw sequencing reads were de-multiplexed using bcl2fastq.

### Host bioinformatics

Mouse reads were processed using Trimmomatic^37^ v0.39 to remove TruSeq3-PE-2 adapter sequences using default parameters and filtering for reads longer than 36 bases after trimming. Reads were aligned to the mm39 genome assembly (GCA_000001635.9) and using Star^38^ v2.7.10b with default parameters. The genome assembly and associated annotation file were acquired from the University of California Santa Cruz Genomics Institute.^39^ Additional alignment metrics were collected using Picard^40^ v2.27.5. Genes with excessive number of zero values (>90%) or low variance across samples were removed from analysis, resulting in a final dataset of 29,579 host genes. Additional details in Supplemental Methods.

### Bacterial bioinformatics

Bacterial reads were trimmed using Skewer v0.2.2^41^ with an adapter sequence of 5’-CTGTCTCTTATACACATCT-3’, a read length range of 50-175 bases, and an end quality threshold of 20. Paired-end reads were reassembled using PairFQ lite^42^ v0.17.0. Reads were then aligned to the 2016 *Mtb* Erdman reference genome assembly using Bowtie2^43^ v2.5.0 with default parameters. Mapped sequences were counted using HtSeq^44^ v1.0 with default parameters. Only genes detected by SEARCH-TB^32^ were included in the analysis (n=3,568 genes). For gene clustering, genes with low variance were further excluded, resulting in a dataset of 3,317 genes. Additional details in Supplemental Methods.

### Clustering genes by longitudinal expression patterns

Expression was normalized using the variance stabilizing transformation in DESeq2,^45^ scaled with the R *scale* function then filtered to select the most variable genes, resulting in a dataset of 18,354 genes. For each gene, the average expression across all samples per time point was calculated. Auto-correlation coefficients (ACF) were calculated to quantify temporal dependencies and characterize dynamic patterns over time, measuring how similar a time series is to itself at different time lags and then using the ACFs, genes were grouped via community clustering based on Euclidean distance, using a K value of 4000 for host genes and 540 for *Mtb* genes (**Fig. S1-2**). Since ACF captures temporal dynamics but not directionality, the resulting clusters contained symmetric patterns. To separate them, we split clusters based on whether gene expression at the second time point was greater than the first time point. This initially resulted in eight clusters for the host and ten clusters for *Mtb* (**Fig. S3**). The *Mtb* clusters were consolidated based on similar temporal expression patterns, yielding six for *Mtb* (**Fig. S4**). Additional details in Supplemental Methods.

### Mouse Pathway Enrichment Analysis

Pathway enrichment analysis was performed for pairwise combinations to evaluate the overrepresentation of differentially expressed genes in Gene Ontology (GO) categories. GO terms were used from Biologic Process, Molecular Function, and Cellular Component ontologies (m5.go.v2023.2.Mm.symbols.gmt). Enrichment analysis was conducted using clusterProfiler^46^ separately for upregulated and downregulated genes. Gene categories with Benjamini-Hochberg^47^ adjusted *p*-values<0.05 were considered significant.

### Differential Gene Expression

We fit negative binomial generalized linear models using the *edgeR* package^48^ using a term for model/timepoint and a term for batch. Likelihood ratio tests assessed differences in expression between different conditions at each timepoint. Genes with a log fold change greater than 0.5 for *Mtb* and 1 for the host and with Benjamini-Hochberg^47^ adjusted *p*-values<0.05 for tests of interest were deemed significant.

### Bacterial Pathway Enrichment Analysis

Functional enrichment analysis was conducted separately for significantly up- and down-regulated gene sets using hypergeometric tests in the *hypeR*^49^ R package for pairwise comparisons to identify overrepresentation in functional categories established by Cole et al.^50^ or curated from the literature (**Table S1**). Categories containing <8 genes were excluded. Benjamini–Hochberg adjusted P-values^51^ <0.05 were considered significant.

### Average expression plots

Normalized expression levels of genes within selected bacterial gene categories were averaged per mouse sample, generating a single value per category per mouse as further described in **Fig. S5**.

### Online analysis tool

Host and pathogen can be explored interactively using an Online Analysis Tool built with Shiny: https://microbialmetrics.org/analysis-tools/.

## RESULTS

### Rapid *Mtb* replication in the LDA and HDA models before the onset of adaptive immunity

Change in CFU and RS ratio were monitored before and after the onset of adaptive immunity (**Fig. 1A**). One day post-infection, there were 2.40 ± 0.07 (SEM) and 3.83 ± 0.26 CFU in the lungs of LDA and HDA mice, respectively. During the innate phase (first 11 days), CFU rose rapidly, increasing 2.59 and 3.02 log in HDA and LDA, respectively (**Fig. 1B**). This represented a doubling time of 22.7 hours (95% CI: 19.8-23.3) in the LDA model and 26.8 hours (95% CI: 24.0-32.8) in the HDA model (**Fig. 1C**) approaching doubling times typically observed at log-phase growth *in vitro*.^52^ Between days 11 and 19, CFU rose more slowly (doubling time ∼50 hours in both models). After day 19, CFU in the LDA stabilized and declined slightly, suggesting that the onset of adaptive immunity resulted in slower *Mtb* replication, increased immune killing or both. In the HDA model, development of signs of distress (i.e., weight loss, hunched posture, lethargy) necessitated humane euthanasia on day 19 so collection of later time points was not possible. As anticipated, CFU was consistently ∼2 log higher in the HDA model than in the LDA model during the innate immune period (**Fig. 1B**).

**Figure 1:**
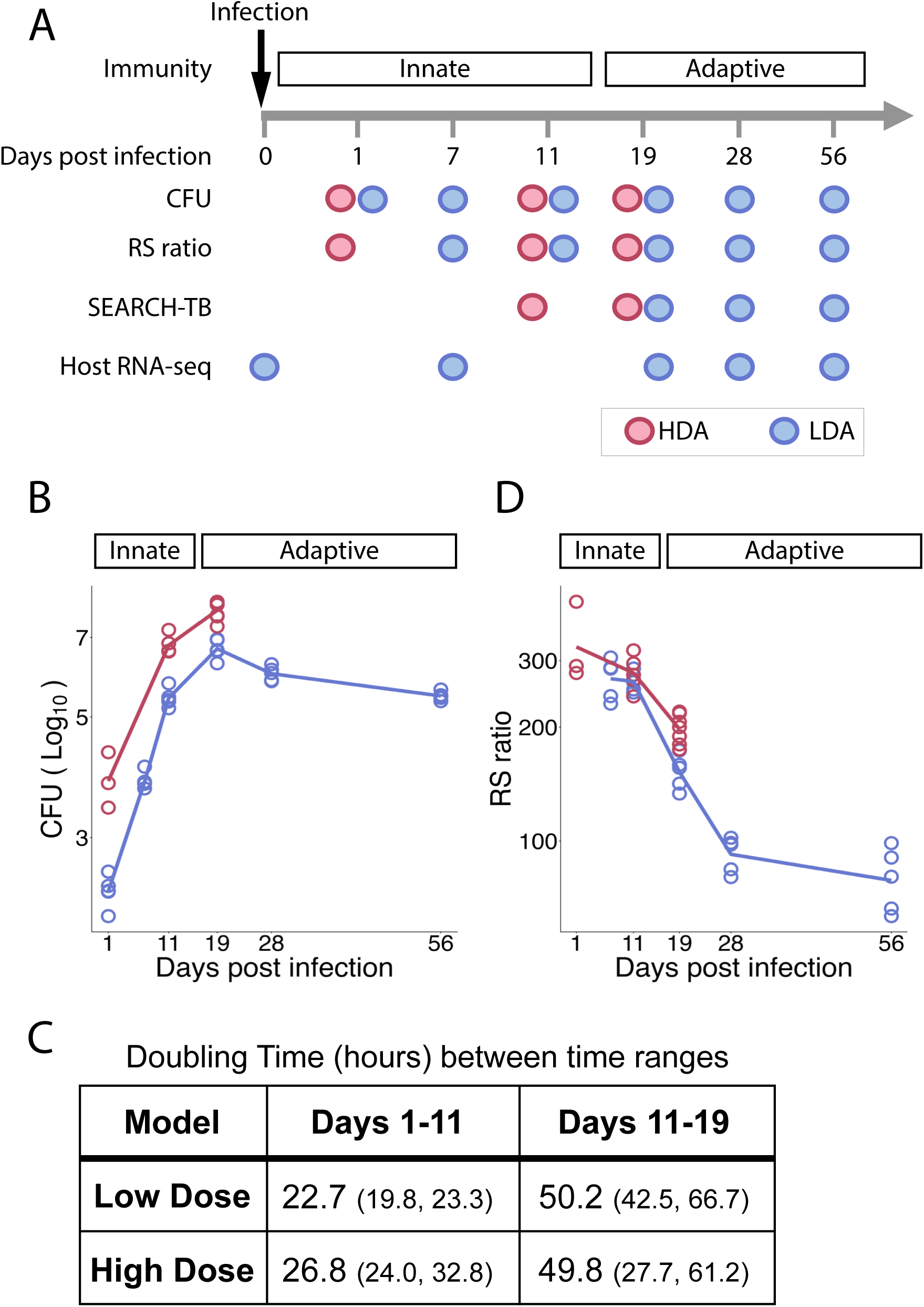
Schematic representation of the experimental design with the bacterial burden and activities. (**A**) Murine subjects were inoculated with either a high-dose aerosol (HDA (infection of ∼10^4^ bacilli), red) or low-dose aerosol (LDA (infection of ∼10^2^ bacilli), blue), and pulmonary tissue was harvested at various time points post-infection. At specified timepoints, a cohort of five mice were sacrificed for specimen collection. Dots denote the implementation of a specified measure on a given day. Analytical measures included colony-forming unit (CFU) enumeration, RS ratio for bacterial activity, SEARCH-TB assay for bacterial RNA gene expression, and Murine RNA sequencing (RNA-seq). RNA-seq analyses was conducted exclusively on LDA samples. RS ratio data was unavailable for day 1 in the LDA model. (**B**) CFU of HDA (red) and LDA (blue) infection over time where data points represent individual murine samples. The median value for each available time point is indicated by a connecting line. (**C**) Doubling time calculated from CFU data for the intervals of days 1-11 and days 11-19. The 95% confidence interval, determined through bootstrap analysis, is presented in parentheses as (lower bound, upper bound). (**D**) RS ratio of HDA (red) and LDA (blue) infections over time where data points represent individual murine samples. The median value for each available time point is indicated by a connecting line.

### Slowdown in *Mtb* rRNA synthesis with onset of adaptive immunity

The RS ratio assay^20^ demonstrated reduction in bacterial rRNA synthesis concordant with the onset of adaptive immunity. In both the HDA and LDA models, the RS ratio was high on days 7 and 11, consistent with active rRNA synthesis during the innate phase (**Fig. 1D**). In the LDA model, the RS ratio declined significantly between day 11 and 19 (*p-value=0.000060*) and again between day 19 and 28 (*p-value=0.0028*) thereafter, suggesting that activated host adaptive immunity constrained *Mtb*, promoting suppressed bacterial rRNA synthesis.

### Longitudinal change in host transcriptome in the LDA model

Interrogation of the mouse lung transcriptome from these same mice allowed us to establish the kinetics of the onset of adaptive immunity. A heatmap identified a distinct shift in host expression between days 11 and 19 (**Fig. 2A**). Unsupervised clustering partitioned expression over time into eight gene clusters, half of which rose between days 11 and 19 and half of which fell during the same period (**Fig. 3B; Fig. S6**). Cluster 1, which did not increase until after day 11, was uniquely enriched for adaptive immune processes (*Adj-P<0.00001*) (**Fig. S7**). Following day 19, expression levels remained stable with subtle fluctuation across day 28 and day 56 (**Fig. 2B**, **Fig. S8**).

**Figure 2:**
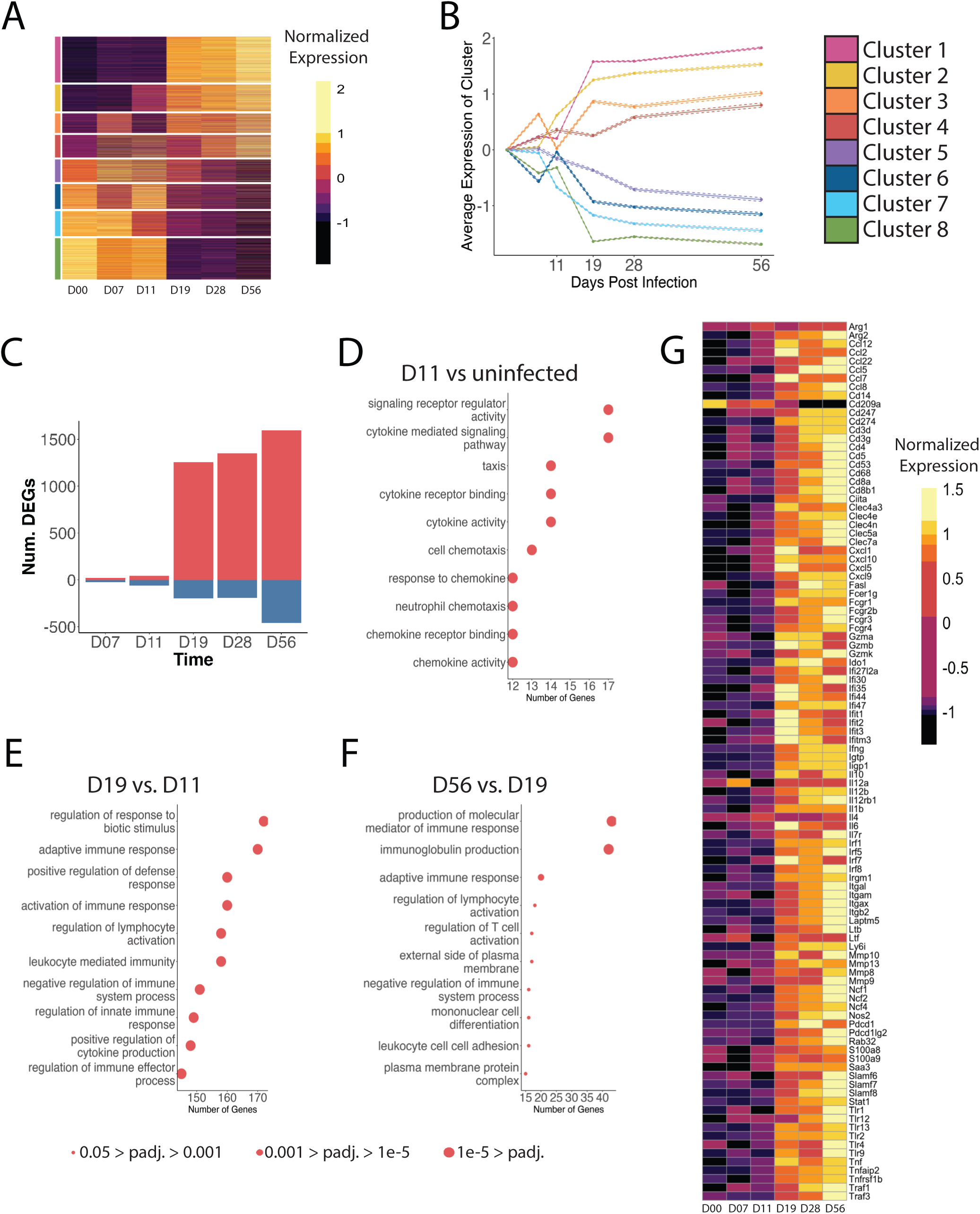
Transcriptional response of Host in the LDA model. (**A**) Heatmap of normalized gene expression of the 18,354 (of 29,579) most variable mouse genes, organized by cluster. Low variance genes were removed from clustering analysis. (**B**) Line plot of average gene expression of mouse genes by cluster. Shaded region indicates 95% confidence interval. (**C**) Quantification of differentially expressed genes (DEGs) at the specified time point compared to uninfected mice (day 0). Red indicates upregulated genes, while blue indicates downregulated genes. (**D-F**) Dot plots illustrating the top 10 gene set enrichments within upregulated genes at day 11 compared to uninfected (day 0) (**D**), day 19 compared to day 11 (**E**), and day 56 compared to day 19 (**F**). Dot size represents adjusted *p*-value by over-representation analysis method with Benjamini–Hochberg correction. (**G**) Heatmap illustrating normalized gene expression of selected immunity-related genes.

**Figure 3:**
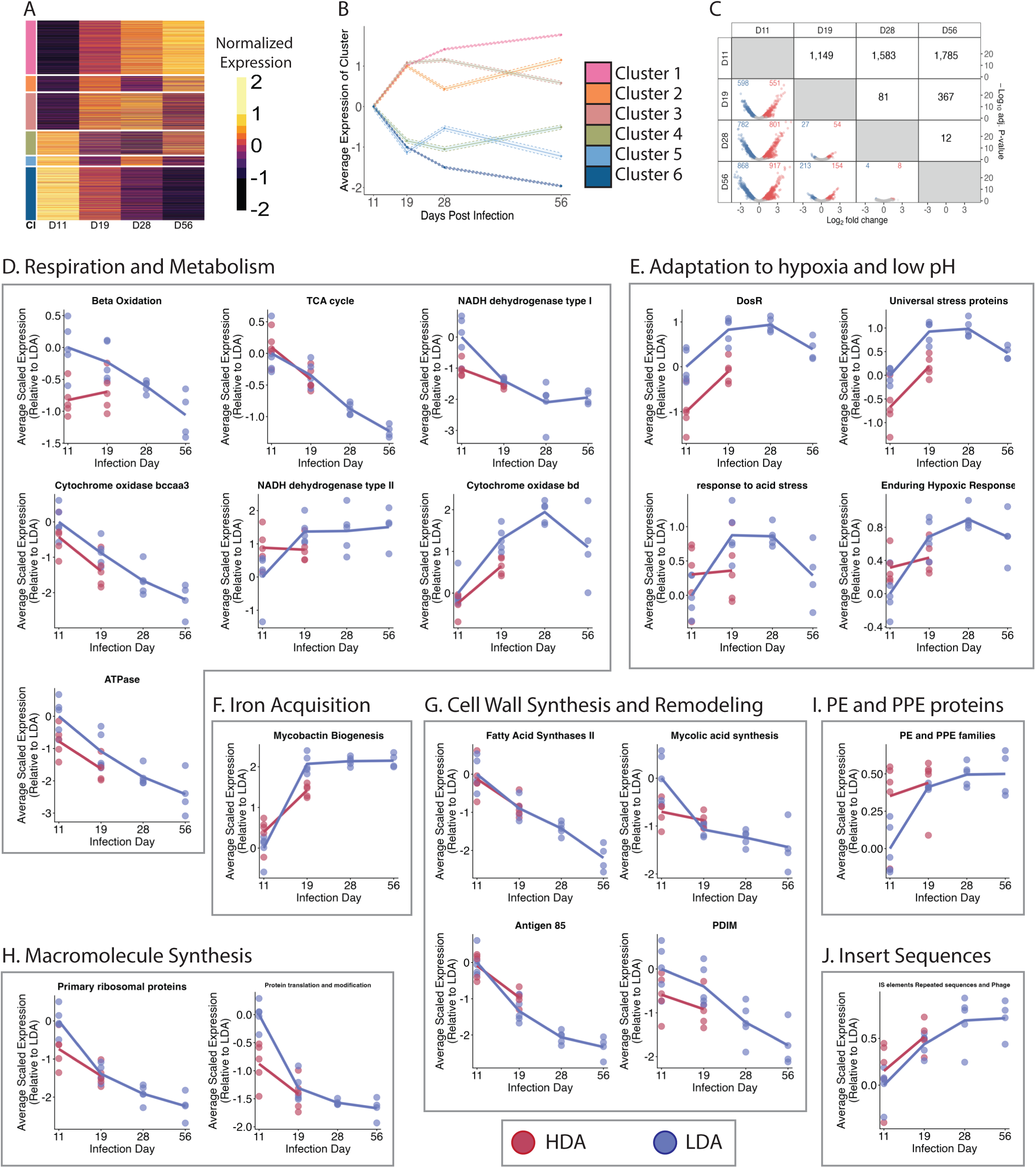
Transcriptional response of *Mtb*. (**A**) Heatmap of normalized gene expression of 3,317 (or 3,568) *Mtb* genes, organized by cluster. Low variance genes were removed from clustering analysis. (**B**) Line plot of average gene expression of *Mtb* genes by cluster. Shaded region indicates 95% confidence interval. (**C**) Matrix of differentially expressed *Mtb* genes between time points of LDA infection. Columns indicate reference and rows indicate comparison group. The upper right quadrant presents the total number of DEGs between time points. The lower left quadrant displays volcano plots where downregulated (blue), and upregulated (red) genes (circles) are plotted by the log fold change and adjusted *p*-value. (**D-J**) Average expression of specified gene set categories per murine sample (dot). A mean trend line was computed by averaging values across all samples at each time point.

Concordant with the above, only 49 and 108 genes were significantly differentially expressed on days 7 and 11 relative to pre-infection, respectively (**Fig. 2C**). Gene set enrichment analysis showed that genes upregulated on day 11 were enriched for cytokine activity (*Adj-P<0.0001*), chemokine activity (*Adj-P<0.00001*), and cytokine-mediated pathways (*Adj-P<0.00001*), consistent with an innate immune response (**Fig. 2D**). By contrast, 1,293 genes were significantly differentially expressed on day 19 compared to day 11 (**Fig. S9**). The gene set upregulated on day 19 relative to day 11 was enriched for adaptive immune response (*Adj-P<0.00001*) and related processes (**Fig. 2E**). Following day 19, transcriptional changes were relatively modest: only 57 and 272 DEGs were detected on days 28 and 56, respectively, compared to day 19 (**Fig. S9**). Most genes differentially expressed on day 19 compared to pre-infection remained altered in the same direction through day 56, indicating that the increased number of DEGs at day 56 (**Fig. 2C**) emerged from genes that were previously not significantly altered in either direction. Transitions from significantly downregulated to upregulated states and vice versa, were rare (**Fig. S10**). Genes upregulated on day 56 compared to day 19 were enriched for immune response mediators and immunoglobulin-related functions (*Adj-P<0.00001* for both) consistent with maturation from an early cellular response to later antibody-mediated response later in infection^53–55^ (**Fig. 2F**).

We further examined individual genes canonically associated with adaptive immunity^26,27^ (**Fig. 2G**). Between days 11 and 19, there was a significant increase in expression of *nos2* and antimycobacterial effectors *cd53*, *cd68*, *laptm5*, *ltf*, *ncf1*, *ncf2*, *ncf4*, *rab10*, *rab20*, and *rab32*.^33^ There was also significantly increased expression of genes for pro-inflammatory chemokines previously shown to be involved in neutrophil, monocyte, and T cell recruitment, including: genes *ccl2*, *ccl5*, *ccl7*, *ccl8*, *ccl12*, *cxcl1*, *cxcl5*, *cxcl9*, and *cxcl10*.^15^ Key adaptive cytokines *ifng* and *tnf* also showed robust induction. For all above genes, the adjusted p-values were *<0.00001*. Fold-changes and exact p-values are shown in **Table S2**.

### Longitudinal change in *Mtb* transcriptome in the LDA model

Clustering bacterial genes (**Fig. 3A-B**) identified a distinct shift in expression between infection days 11 and 19 corresponding to transcriptional changes observed in the host. Unsupervised clustering partitioned expression over time into six gene clusters, half of which rose between days 11 and 19 and half of which fell during the same period (**Fig. 3B, Fig. S11**). Qualitatively, all six gene clusters indicated a sharp change in expression between days 11 and 19 concordant with the onset of adaptive immunity observed in the murine host, with more modest changes thereafter. Categorical enrichment of the six clusters is summarized in **Table S3**. The pattern of rapid transcriptional change between days 11 and 19 followed by transcriptional stabilization was supported quantitatively by the number of genes that were significantly differentially expressed between pairs of timepoints. The greatest transcriptional change (1,149 DEGs) occurred between days 11 and 19. By contrast, only 81 DEGs were observed at day 28 compared to day 19 and only 12 DEGs at day 56 compared to day 28 (**Fig. 3C**). In the following sections, we evaluated longitudinal changes in selected *Mtb* physiologic processes. Additional processes can be evaluated via the accompanying online analysis tool: https://microbialmetrics.org/analysis-tools/.

### Decreased Respiration and Metabolism

Progressive decrease in bacterial aerobic metabolism and oxidative phosphorylation during the onset of adaptive immunity was evidenced by a progressive decrease relative to day 11 in expression of *Mtb* genes involved in beta oxidation (*Adj-P=0.047* on day 28) and the TCA cycle (*Adj-P=0.047* on day 19). Additionally, there was a progressive decrease in primary proton-pumping NADH dehydrogenase Type I (*Adj-P=0.00054* on day 19), and the primary terminal oxidase cytochrome *bcc/aa3* supercomplex (*Adj-P=0.025* on day 19) that *Mtb* uses for efficient aerobic respiration in oxygen-replete growth conditions (**Fig. 3D**). Concordantly, a metabolic shift was indicated by increased expression of the non-proton-pumping NADH dehydrogenase Type II (*Adj-P=0.0077* on day 19) and the less efficient cytochrome *bd* oxidase (*Adj-P=0.0014* on day 19) (**Fig. 3D**) used during hypoxic or acid stress.^56^ Concordant with decreased energy production, expression of ATP synthase genes decreased progressively (*Adj-P=0.0076* on day 19) (**Fig. 3D**).

### Adaptation to hypoxia and low pH

Expression of DosR genes, a regulon canonically transiently induced by stressors encountered within activated macrophages including hypoxia, nitric oxide, and carbon monoxide^57^ was induced relative to day 11 (*Adj-P=0.0031* on day 19) then returned to baseline on day 56 (**Fig. 3E**). Similarly, expression of genes for Universal Stress Proteins (USPs), which respond to diverse environmental stresses such as hypoxia, low pH, and NO exposure^58^ rose transiently (*Adj-P=0.00020* on day 19) and then returned to baseline on day 56 (**Fig. 3E**), suggesting a role in early adaptation to environmental stress. Expression of genes associated with acid stress rose transiently (*Adj-P=0.013* on day 19), then returned to baseline on day 56 (**Fig. 3E**). Expression of a set of genes described as the enduring hypoxic response^57^ rose significantly (*Adj-P=0.0013* on day 19) and remained high thereafter (**Fig. 3E**).

### Increased expression of genes related to iron acquisition

Relative to day 11, expression of genes involved in synthesizing mycobactin, a siderophore essential for iron sequestering from the environment, were induced (*Adj-P<0.00001* on day 19) and remained elevated through the chronic phase (**Fig. 3F**), suggesting *Mtb* adaptation to iron-limited conditions and the need to repair iron-containing enzymes damaged by ROS/RNS from immunity.^59^

### Decreased cell wall synthesis and remodeling

Relative to day 11, expression of genes for key cell wall constituents and processes decreased, including the fatty acid synthase II (FAS-II) that synthesizes mycolic acid precursors (*Adj-P=0.0070* on day 19), mycolic acid modification (*Adj-P=0.00074* on day 19), and Antigen 85 mycolytransferase that transfers mycolic acids to the cell surface (**Fig. 3G**) (*Adj-P=0.000015* on day 19). Additionally, there was decreased expression of phthiocerol dimycocerosate (PDIM) cell surface glycolipids genes (*Adj-P=0.0076* on day 28) (**Fig. 3G**) and peptidoglycans that form the structure of the *Mtb* cell wall (*Adj-P=0.00054* on day 19) (**Fig. S12**).

#### Decreased macromolecule synthesis

Relative to day 11, there was progressively decreased expression of genes encoding primary ribosomal proteins (*Adj-P=0.000047* on day 19) (**Fig. 3H**) and involved in ribosomal modification and maturation (*Adj-P=0.0013* on day 19) (**Fig. S12**). In contrast, the four alternative C-ribosomal protein paralogs with a role in stress-induced ribosomal remodeling had gradual increased expression (*Adj-P=0.0062* on day 28) (**Fig. S12)**. Concordantly, genes for protein translation and modification decreased during the onset of adaptive immunity (*Adj-P<0.00001* on day 19) (**Fig. 3H**). Finally, the broad category of macromolecule synthesis and processing decreased (*Adj-P=0.000045* on day 19) (**Fig. S12)**, indicating global slowing of bacterial activity.

### Regulation of growth: Sigma factors

As illustrated in **Fig. S13**, the onset of adaptive immunity was associated with markedly altered expression of the 13 sigma factors that bind RNA polymerase to regulate gene expression. Relative to day 11, expression of *sigA*, which codes for the primary ‘housekeeping’ sigma factor necessary for growth, was profoundly and consistently suppressed (*Adj-P<0.00001* on days 19-56). By contrast, certain stress-associated alternative sigma factors associated with growth arrest (*sigE, sigF, sigH*) were consistently significantly induced on days 19-56, relative to day 11 (p-values shown in **Table S4**).

### Post-transcriptional regulation of stress response: Toxins

As illustrated in **Fig. S14**, the onset of adaptive immunity was associated with markedly altered expression of genes for toxins that act post-transcriptionally to reprogram *Mtb* in response to stress, with certain toxin genes (*vapC18, higB, Rv3749c, Rv2019, mazF6 and relG*) consistently, and significantly, induced relative to day 11 with others consistently significantly suppressed (p-values shown in **Table S5**).

#### PE and PPE proteins and Insertion Sequences

Relative to day 11, there was increased expression of genes for PE and PPE proteins that play diverse roles in immune evasion and survival^60^ (*Adj-P=0.0014* on day 19) (**Fig. 3I**) and insertion sequences that are recognized as a component of adaptation to host stress (*Adj-P=0.013* on day 19) (**Fig. 3J**).^61^

### Concordance of host and *Mtb* transcription

Since the most pronounced shifts in gene expression for both host and pathogen occurred between infection days 11 and 19, we juxtaposed change in five host immune-related processes with four key *Mtb* processes (**Fig. 4**). This highlighted that increased expression of host genes related to activated T-cell proliferation, antigen processing and presentation, B-cell activation, macrophage activation, and neutrophil activation coincided with increased expression of bacterial DosR regulon and mycobactin synthesis genes. Conversely, expression of *Mtb* aerobic respiration and primary ribosomal protein synthesis genes was inversely correlated with the immune processes. The accompanying online analysis tool (https://microbialmetrics.org/analysis-tools/) enables expression of any host gene category to be juxtaposed with expression of any *Mtb* gene category.

**Figure 4:**
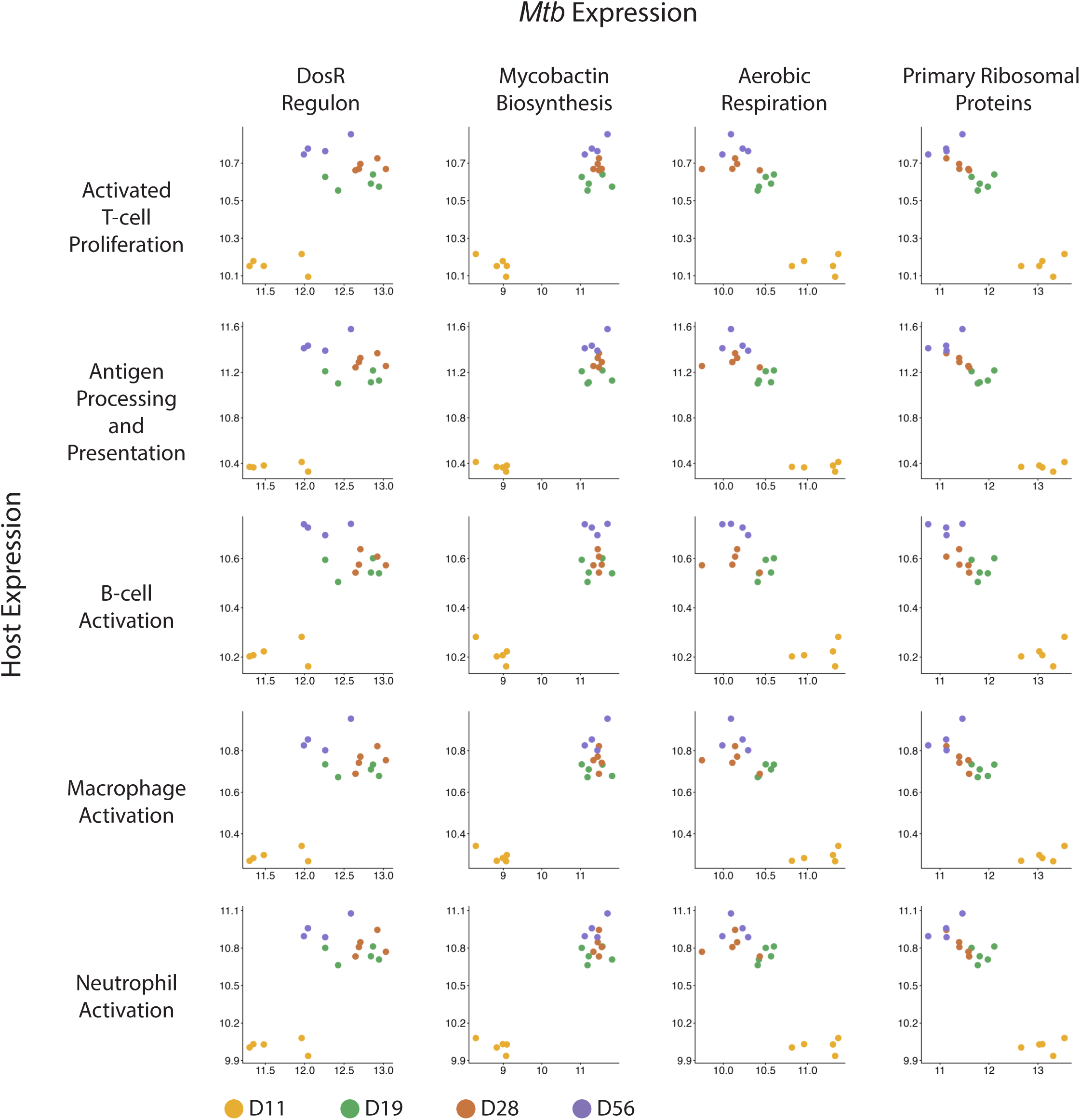
The transcriptional interaction between Host and *Mtb* in the LDA model. Mean of the normalized expression level of *Mtb* gene category (x-axis) in relation to the mean of the normalized expression level of selected host gene groups (y-axis). Each data point represents a mouse sacrificed on the day indicated by color.

### Change in *Mtb* transcriptome in the HDA model

In the HDA model, only two time points were evaluable because disease progression necessitated humane euthanasia. Between days 11 and 19, the number of genes differentially expressed (N=171) (**Fig. S15**) and average log fold change (0.35x) were lower than observed between these timepoints in the LDA model (N=1,149; 0.61x) (**Fig. 3C**). Of the changes that did occur, many of the same gene categories were significantly different in the HDA model as were observed in the LDA model during the same interval after *Mtb* challenge (**Fig. 3D-J**).

### Baseline *Mtb* physiology in the HDA (day 11) versus LDA (day 28) models

Finally, we compared the *Mtb* transcriptome at the closest pre-treatment starting points of the HDA and LDA models that are a mainstay of contemporary preclinical evaluation (days 11 and 28, respectively). This revealed 822 differentially expressed genes.

When comparing *Mtb* physiology between HDA day 11 and LDA day 28, we observed clear contrasts that mirror the shifts seen within the LDA model between day 11 and day 19, suggesting that the onset of adaptive immunity was the primary driver of physiological differences between HDA day 11 and LDA day 28. Specifically, transcriptional profiles of HDA at day 11 reflected a physiologically active state, marked by elevated expression of genes involved in protein synthesis and ATP-producing aerobic respiration. In contrast, LDA day 28 displayed a more stressed, immune-constrained phenotype, characterized by upregulation of the DosR regulon, stress response pathways, and iron sequestration mechanisms (**Table 1**). Overall, the shared physiological differences across both comparisons emphasize the distinct, stress-adapted nature of *Mtb* at day 28 following LDA compared to the *Mtb* phenotype present during the innate immune phase at day 11 following HDA.

**Table 1:** Categorical enrichment of *Mtb* gene categories at the baseline of the LDA model (infection day 28) relative to the baseline of the HDA model (infection day 11)

| Grouping | Gene Category | Direction in LDA relative to HDA | Adjusted p-value |
| --- | --- | --- | --- |
| Hypoxia and low pH | DosR regulon | Up | <0.00001 |
|  | Enduring Hypoxic Response | Up | <0.00001 |
|  | Oxidative Stress | Up | <0.00001 |
|  | Universal Stress Proteins | Up | <0.00001 |
| Nutrient acquisition | Mycobactin Biosynthesis | Up | <0.00001 |
|  | EXS-3 | Up | <0.00001 |
|  | Zur regulon | Up | 0.000024 |
| Protein and macromolecule synthesis | Primary Ribosomal Proteins | Down | <0.00001 |
|  | Synthesis and modification of macromolecules | Down | <0.00001 |
|  | Protein translation and modification | Down | 0.023 |
| Metabolism | TCA cycle | Down | 0.00011 |
|  | ATP Synthase | Down | 0.023 |

## DISCUSSION

We used novel molecular assays of pathogen health to characterize bacterial physiology at the baseline stage (pre-treatment) in the BALB/c mouse based on two common preclinical murine TB efficacy models that are central to contemporary TB drug development. Analyses of *Mtb* and host expression demonstrated that the onset of adaptive immunity coincided with transformation in *Mtb* physiology from a rapidly growing, metabolically active phenotype with ongoing macromolecule synthesis to a stressed and metabolically inactive phenotype with diminished macromolecule synthesis. The baseline difference between *Mtb* phenotypes in the HDA and LDA models may explain why many drugs exhibit greater bactericidal activity in the HDA model. Broadly, these results highlight the crucial role of adaptive immunity in shaping *Mtb* physiology which in turn influences drug efficacy.

Our results confirmed that, in the BALB/c mouse, innate immunity fails to control *Mtb,* allowing exponential growth similar to log-phase *in vitro* replication. During unconstrained bacterial growth in the innate immune phase, *Mtb* had a metabolically active phenotype with high macromolecule and cell wall synthesis and low expression of stress responses. Consistent with previous studies,^15,33–35^ host transcriptional changes during the innate phase were modest, limited primarily to cytokine and chemokine induction without evidence of effector cell activation. The onset of adaptive immunity was accompanied by massive transcriptional change in both host and pathogen. When host expression indicated immune effector cell activation (in our experiments at day 19 or later), *Mtb* expression shifted to a stressed phenotype that had decreased macromolecule and cell wall synthesis. Bacterial metabolism appeared to slow and shift to alternative, less-efficient respiration.

There was increased expression of canonical markers of the effector mechanisms used by activated macrophages^62^ and corresponding up-regulation of canonical bacterial adaptive responses.^63,64^ For example, concordant with increasing host expression of *Nos2* that encodes inducible nitric oxide synthase and *Hif1A* that encodes Hypoxia-Inducible Factor 1 Subunit Alpha, *Mtb* sharply increased expression of the DosR regulon that responds to hypoxia and nitric oxide. The hypothesis that DosR may be an indirect readout of adaptive immunity is consistent with our previous observation that the DosR regulon had lower expression in the sputum of TB patients with compromised adaptive immunity due to advanced AIDS than in immunocompetent patients.^62,65^ Similarly, acidification of the phagolysosome is recognized as a central anti-microbial mechanism of activated macrophages^66,67^ and we observed a corresponding adaptive increase in expression of pH responsive *Mtb* genes. Finally, activated macrophages sequester iron^68^ to deprive *Mtb* of an enzymatic co-factor essential for growth and survival. During the onset of adaptive immunity, we observed a corresponding increase in expression of *Mtb* siderophore genes that scavenge iron.

The transition in *Mtb* physiology that occurred with the onset of adaptive immunity was broadly similar in the HDA and LDA models but of greater magnitude in the LDA model (*i.e.,* more differentially expressed genes and larger fold changes). We speculate that in the HDA model overwhelming infection might have imposed less intense and less effective immune stress on a “per bacillus basis” than in the LDA model, resulting in a diminished bacterial adaptive response.

Our results highlight that the HDA subacute TB efficacy model that is the mainstay of TB drug evaluation should be understood as a “mixed model” in which drug exposure spans both the innate and adaptive immune phases. During the first days of the HDA treatment model, drugs act on the rapidly growing, metabolically active *Mtb* phenotype that innate immunity fails to contain. Within the first week, adaptive immune pressure is added. Our results here with the LDA model demonstrate that, if mice are not overwhelmed by exponential *Mtb* growth in the innate phase, adaptive immunity alone is sufficient to contain *Mtb*. By reducing bacterial burden and rescuing HDA mice from otherwise lethal overwhelming infection, drug treatment “buys time” for the onset of adaptive immunity. The idea that adaptive immune pressure rapidly becomes a factor in the HDA model is supported by our separate recent experiment that treated BALB/c HDA mice with a 2-week course of isoniazid, rifampin, pyrazinamide and ethambutol then stopped treatment and monitored recovery in *Mtb* physiology.^69^ We found that even one month after treatment discontinuation, *Mtb* did not rebound with rapid growth, but instead retained a quiescent, stressed phenotype similar to the phenotype we observed here in the chronic LDA model. We hypothesize that brief drug treatment that reduced *Mtb* burden and allowed for the onset of adaptive immunity essentially “converted” the HDA model into the LDA model.

These observations are important for several reasons. The LDA and HDA models have historically been viewed as fundamentally different^5^ because LDA begins with a slowly replicating immune-constrained *Mtb* phenotype and HDA begins with a rapidly replicating unconstrained *Mtb* phenotype. This observation leads to the critique that the HDA model does not recapitulate slow-growing *Mtb* phenotypes found in humans. However, this study and our previous experiments suggest that the period between treatment onset in the HDA model and the onset of adaptive immune pressure is actually quite short (∼1 week). Since adaptive immunity is present for the subsequent months, the HDA relapse model does measure sterilizing efficacy against a chronically immune-constrained intra-cellular *Mtb* population. Second, by revealing the profound effect of adaptive immunity on *Mtb* physiology, our results are a reminder of the need to consider the contribution of immunity to drug efficacy. During treatment, there is a three-way interaction between the pathogen, the host and drug pressure. Our current work evaluated only the interaction between pathogen and host but lays the foundation for future evaluation of the effect of drugs on host-pathogen interaction.

Host-pathogen interactions in murine TB models have traditionally been evaluated by measuring CFU which estimates the burden of *Mtb* capable of growth on agar. CFU provides no information on bacterial physiologic processes. Our work highlights the extraordinarily granular information that can be gained using molecular assays of pathogen health such as SEARCH-TB and RS ratio. Here, we used these tools to evaluate the role of adaptive immunity in shaping baseline *Mtb* phenotypes in mice by contrasting two commonly employed preclinical TB efficacy models. Our long-term objective is to advance these molecular tools as practical advanced pharmacodynamic markers because they reveal drug effects and interactions that are imperceivable based on CFU alone.^32,69–71^ In particular, these results showed that the RS ratio can serve as a functional readout of the onset of adaptive immunity. Future murine studies will use the RS ratio to evaluate the effect of vaccines on the timing and effectiveness of adaptive immunity in limiting bacterial rRNA synthesis.

This study has several limitations. Both host RNA-seq and SEARCH-TB were performed on lung homogenate. The host transcriptome described here therefore is the average signal across diverse host cell types. In future studies, profiling specific cell types or single cell profiling would enable more granular understanding of host responses. For the bacterium, evaluation of lung homogenate is appropriate because the BALB/c mouse develops diffuse, uniform lung infiltrates in which *Mtb* is nearly entirely intracellular (*i.e.,* limited lesional diversity). In the future, a similar longitudinal study could be performed in a model such as the C3HeB/FeJ mouse that develops complex heterogenous intra- and inter-host pulmonary pathology in response to virulent *Mtb* infection. Finally, we recognize that temporally concordant change in host and pathogen transcription alone does not establish that host pressure caused the transformation in bacterial physiology. However, the patterns of change in host and pathogen transcription are consistent with established mechanisms of host-*Mtb* interaction.

Overall, our findings highlight key differences in the physiological state of *Mtb* during the typical treatment initiation period across two widely used preclinical drug evaluation models. We found that the onset of adaptive immunity was associated with a marked change in *Mtb* phenotype that likely underlies reduced drug activity and efficacy historically observed in the LDA model.^2,72^ However, our results suggest that, in the HDA model, the onset of adaptive immunity begins shortly after the traditional treatment starting point so the HDA also tests an immune-constrained *Mtb* phenotype. Broadly, these experiments emphasize the profound effect of adaptive immunity on *Mtb* phenotypes and the importance of considering the interactions between drug, pathogen, and host.

## ACKNOWLEDGEMENTS

We acknowledge the staff of the Laboratory Animal Resources at Colorado State University for their animal care.

## FUNDING

NW, MV and GR acknowledge funding from NIH UM1 AI179699.

